# The native human glomerulus features a slit diaphragm resembling a densely interwoven fishnet

**DOI:** 10.1101/2025.09.24.678239

**Authors:** Deborah Moser, Alexandra N. Birtasu, Lilli Skär, Pauline Roth, Lisa Rehm, Mike Wenzel, Jens Köllermann, Mbuso S. Mantanya, Felix K.H. Chun, Margot P. Scheffer, Achilleas S. Frangakis

## Abstract

The slit diaphragm (SD), a core element of the glomerular filtration barrier, is essential for renal filtration and disrupted in all forms of glomerulopathy. Using cryo-electron tomography of human kidney tissue, we resolved the near-native SD architecture at unprecedented resolution. The SD comprises crisscrossing strands intersecting at ∼90°, forming a fishnet-like lattice across the ∼44 nm space between podocyte foot processes. An atomic model based on the Nephrin–Neph1 heterodimer reveals ∼9 nm spacing in humans, compared with 12.3 nm in mice and 15 nm in Drosophila. Our data establish the SD as a conserved fishnet-like assembly from invertebrates to mammals, with the tighter human lattice likely conferring enhanced permselectivity.

## Main Text

A comprehensive understanding of the molecular architecture of the slit diaphragm (SD), a critical component of the glomerular filtration barrier, is essential in elucidating renal filtration physiology and the mechanisms underlying kidney disease (1, 2). Disruption of the SD is a hallmark of all forms of glomerulopathy, regardless of whether the cause is genetic, immunological, metabolic, or vascular. Our recent work demonstrated that the SD architecture in mice and *Drosophila* resembles a fishnet, with species-specific structural adaptations (3, 4). However, the architecture of the human SD – which is central to human glomerular disease – remained unresolved. Here, we report the near-native *in situ* architecture of the human SD visualized at an unprecedented resolution through cryo-electron tomography (cryo-ET) of human kidney tissue.

Renal tissue was obtained from an adult patient undergoing a right nephrectomy for renal cell carcinoma (Figure 1a). For cryo-ET, non-tumorous renal cortex was sampled from a region distal to the tumor margin, representative of healthy kidney tissue (Figure 1b). From the healthy kidney tissue, we successfully extracted 18 glomeruli of ∼150 µm in diameter. Processing tissue of this size for cryo-ET, particularly native human tissue, was previously unattainable due to technical limitations. We overcame these challenges by implementing a dedicated preparation pipeline combining high-pressure freezing (without the use of any chemicals or heavy metal staining), focused-ion beam milling and cryo-ET (Supplementary figure S1).

**Figure 1:**
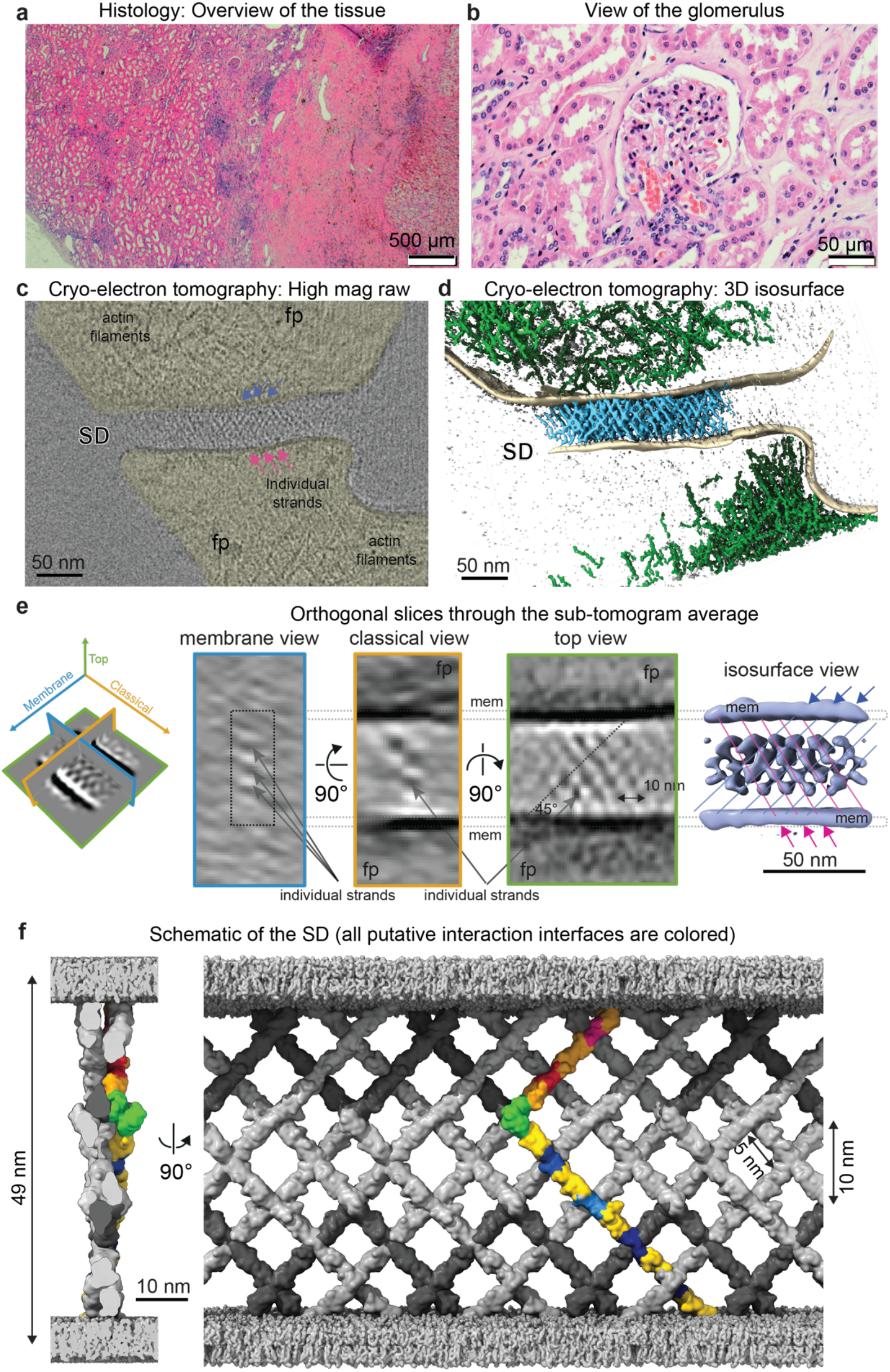
Architecture of the human glomerular filtration system. **(a)** Overview image of a conventional histology of a kidney biopsy from a nephrectomized human kidney showing the overall tissue morphology (hematoxylin stains nuclei dark purple/blue and eosin stains cytoplasm and extracellular matrix pink). On the left, the healthy tissue from the renal cortex can be recognized. On the right, the cancerous tissue is visible. **(b)** Higher-magnification light microscopy image showing the normal architecture of a glomerulus and surrounding non-tumorous tissue. **(c)** Computational slice (1 nm thick) through a raw cryo-electron tomogram (not processed and not denoised) of the filtration system. Foot processes (yellow) contain branching actin filaments and actin bundles. The top view (en face) of the SD shows criss-crossing strands forming a fishnet; strands spanning the membrane from top right to bottom left of the image are indicated by blue arrows, while strands extending from top left to bottom right are indicated by pink arrows. The arrows are spaced 9 nm apart. **(d)** Isosurface representation of the denoised and segmented cryo-electron tomogram (in c) displaying the SD (in blue) from the top-view spanning the two plasma membranes (in beige) of the foot processes. Large areas of the foot processes are filled with branched actin (in green). **(e)** Three orthogonal views (membrane view (blue frame), classical view (orange frame) and top view (green frame)) of computational sections through the sub-tomogram average of 62 SD segments. Criss-crossing densities span the plasma membranes (indicated with “mem”) of foot processes. Membrane and classical views show individual strand sections (dots), and top and isosurface views reveal a fishnet pattern with the strands crossing at ∼90°. The isosurface view of the sub-tomogram average shows the fishnet pattern with internal structural holes. The diagonal light blue and pink lines indicate crossing strands, highlighting the fishnet pattern. The lines are spaced 9 nm apart. **(f)** Idealized molecular model of the human SD shown in two orthogonal views. The Nephrin-Neph1 heterodimer (AlphaFold) was used as a unit-cell and placed in one-dimensional crystal packing. One heterodimer is highlighted in gold with the putative interfaces shown in various colors. All other heterodimers are shown in grey (Nephrin in light grey and Neph1 in dark grey). Each heterodimer is predicted to interact with six other heterodimers. The spacing between the heterodimers is 10 nm.

We successfully prepared thin cryo-lamellae from two glomeruli and recorded 5 tilt-series in which equidistant strands (∼9 nm apart) spanning the foot processes can be discerned (Supplementary figure S2). In the raw tomograms, the SD spans adjacent podocyte foot processes, forming a fine, web-like mesh of crisscrossing strands across the extracellular space, resembling a fishnet (Figure 1c, Supplementary figure S3). The underlying actin cytoskeleton is also clearly resolved, appearing bundled along the axis of the foot processes and increasingly branched and loosely organized near the plasma membranes (Figure 1d). From the tomograms, 62 SD segments were computationally extracted and averaged, yielding a three-dimensional density map of the human SD at a resolution of ∼4.5 nm (Figure 1e). The map reveals crisscrossing strands intersecting at ∼90 degrees, forming a fishnet-like pattern spanning the ∼44 nm extracellular space between foot processes (center-to-center of plasma membrane, ∼49 nm). Using a hybrid of available crystallographic structures (ID: 4OFY) (5) and AlphaFold predictions (IDs: AF-Q9QZS7-F1, AF-Q80W68-F1), we built an atomic model of the Nephrin-Neph1 heterodimer, the main constituents of the SD that belong to the immunoglobulin (Ig) superfamily cell-adhesion molecules and consist of 10 and 5 Ig-domains, respectively (6). The heterodimers were then assembled into a model of the human SD that reflected the densities of the cryo-ET map (Figure 1f, Supplementary figure S4a).

The SD is more densely woven in humans than in mice (3) or *Drosophila* (4), with individual strands crisscrossing to form a tight mesh (Figure 1f). We showed in mice that these strands correspond to Nephrin–Neph1 heterodimers (3, 5). The heterodimers in humans are spaced ∼9 nm apart, which is closer than in mice (12.3 nm) or Drosophila (15 nm), resulting to seven crossing points per heterodimer compared to four crossing points in mice and three in *Drosophila*. This tighter organization in the human SD produces smaller structural holes of ∼5 × 5 nm versus ∼7 × 7 nm in mice and ∼12 × 12 nm in *Drosophila* (Figure 1f). The smaller structural holes likely enhance permselectivity (Supplementary figure S4b). In mice, each heterodimer crossing point confers specific biophysical properties (such as electrostatic interactions) that facilitate the interaction between individual Ig-domains and contribute both to the stability and assembly of the SD. Specifically, when a molecule is removed from the SD the unoccupied interaction sites at the vacant crossing points will guide the incorporation of the next molecule, which extends into the extracellular space to pair with its partner from the opposing cell. The additional interaction points in the human SD suggest greater heterodimer stability and a more robust filtration barrier, reflecting a functional specialization and adaptation of the human glomerular filtration barrier.

In conclusion, we show that the human SD forms a finely webbed fishnet, an architecture that is evolutionarily conserved from *Drosophila* to mouse to human.The tighter organization in the human SD likely has immediate consequences for the permselectivity, which we hypothesize to be higher than in mice or *Drosophila*, as the passage of filtrates is more restricted (Supplementary figure S4b) (7). Importantly, our ability to analyze native human tissue (without chemical fixation or staining) by cryo-ET at a nanometer resolution enables physiological studies and mechanistic insight and opens new opportunities for the *in situ* investigation of disease mechanisms and therapeutic development.

### Caveats of our study

include restriction to a single patient, as we could not obtain an additional nephrectomized human kidney from a different patient in a reasonable time frame. While the fishnet architecture of the SD is unambiguous, the generation of the human SD model relies on the much better resolved studies of the mouse and *Drosophila* SD. Thus, while all future models of the human SD will look like fishnets, insights into the exact molecular arrangement will improve with more data.

## Supplemental Data

### Materials & Methods

#### Isolation of the human glomeruli

Isolation of glomeruli from nephrectomized human kidney tissue was performed according to an adapted isolation protocol (6). All steps were performed at 4°C or on ice. In brief, a small tissue piece was harvested from nephrectomized human kidney tissue and minced into small pieces (∼1 mm^3^) in Hanks’ Balanced Salt Solution (HBSS, #2323615, Gibco, Thermo Fisher Scientific Inc., Waltham, MA, USA). Minced tissue pieces were gently pressed with a wooden tongue depressor (Glaswarenfabrik Karl Hecht, Sondheim vor der Rhön, Germany), first through an HBSS pre-wetted 150 µm cell strainer and then a 100 µm cell strainer (PluriStrainer, PluriSelect Life Sciences, Leipzig, Germany). The retained sample was rinsed into a centrifuge tube and centrifuged at 115 *x g* for 5 min at 4°C. The pellet was resuspended in 100 µl CellBrite Steady Membrane Stain 550 (#30107, Biotium, Fremont, CA, USA) in HBSS (1:1000) and incubated for 25 min, followed by vitrification by high-pressure freezing.

#### Sample vitrification by high-pressure freezing

Vitrification of the sample was based on a modified preparation protocol (7). Planchets (type B, Wohlwend, Sennwald, Switzerland) were coated with cera alba (1% w/v in diethyl ether) prior to freezing. Formvar-coated grids (copper/palladium, 75-mesh parallel single bar; G2018D, Plano, Wetzlar, Germany) were placed on a planchet, and 3 μl of isolated glomeruli and tubular fragments mixed with Ficoll PM 400 (20% v/v in HBSS) (Sigma-Aldrich, St. Louis, MO, USA) were applied before the sandwich was completed by adding a second planchet. The sample was then high-pressure frozen using an HPM-010 (Abra Fluid, Widnau, Switzerland). Samples were kept at a temperature low enough to prevent ice crystallization and to guarantee that both the sample and the ice remained in a vitreous state.

#### Cryogenic confocal laser scanning microscopy

High-pressure frozen grids were imaged at -196°C by confocal laser scanning microscopy (cryo-CLSM) (CMS196 cryo-stage, Linkam, Salfords, United Kingdom; LSM700, Carl Zeiss, Jena, Germany). Optical configurations were adjusted to capture the autofluorescence of the glomeruli in the green channel (excitation wavelength of 488 nm), the membrane stain in the red channel (excitation wavelength of 555 nm), and the reflection from the grids in the far-red channel (excitation wavelength of 639 nm). Images were acquired with a 5x/ NA 0.16 objective. Data acquisition was performed with Zeiss ZEN 2009 (blue) v2.1.

#### Cryo-focused ion beam milling

The high-pressure-frozen electron microscopy (EM) grids were clipped into cryo-focused ion beam (FIB) autogrids (#1205101, Thermo Fisher Scientific Inc.) and loaded into an EM grid holder under liquid nitrogen using an EM vacuum cryo-transfer system VCT500 loading station (Leica Microsystems, Wetzlar, Germany). The sample was transferred to an ACE600 high-vacuum sputter coater (Leica Microsystems) for platinum coating (8 nm) using an EM VCT500 transfer shuttle via a VCT dock (Leica Microsystems). The sample holder was subsequently transferred into a FIB scanning electron microscope (SEM), i.e. a dual-beam instrument (Helios 600i Nanolab, Thermo Fisher Scientific Inc.) that is equipped with a band-cooled cryo-stage equilibrated at - 163°C (Leica Microsystems). The EM grids were imaged using the SEM (3 kV, 0.21 pA) and the FIB source (gallium ion source, 30 kV, 18 pA). An overview SEM image was acquired at 90° incident angle. An organometallic platinum layer of a few microns thickness was deposited with the gas injection system. Regions of interest were identified by correlating the cryo-CLSM images with the SEM images using Bigwarp. Localized positions were milled by applying a specific stress-relief gap for waffled grids. The net incident angle varied between 23° and 28°. The following FIB currents were used: 9.4 – 45 nA (trenching), 2.5 nA stepwise down to 83 pA (thinning), 240 pA (notch milling), and 33 pA (polishing). During thinning steps, a second platinum sputtering was done (8 nm), followed by deposition of an organometallic platinum layer that was a few microns thick.

#### Cryogenic transmission electron microscopy imaging and cryo-electron tomography

Cryo-FIB milled lamellae (2 glomeruli in total) were imaged using a Titan Krios cryogenic transmission electron microscope (cryo-TEM; Thermo Fisher Scientific Inc.) operating at 300 kV in nanoprobe EFTEM mode, equipped with an X-FEG field emission-gun, a GIF Quantum post-column energy filter operating in zero-loss mode and a K3 direct electron detector (Gatan Inc., Pleasanton, CA, USA). Low-magnification images (×11500) of the lamellae were acquired at an angle between 23° and 28° to match the lamella pre-tilt angle induced by FIB milling (calibrated pixel size 1.642 nm/pix, –50 µm defocus).

Tilt series were recorded at a nominal magnification of ×33,000 (calibrated pixel size 1.34 Å/pix) in super-resolution and dose fractionation mode. The cumulative total dose per tomogram was between 130 e− Å-2, and 150 e− Å-2, and the tilt series covered an angular range from −66° to +66° in reference to the lamella pre-tilt, with an angular increment of 2° to 3° and the nominal defocus set to –5 µm. The complete preparation and recording pipeline are presented in Supplementary figure S1.

#### Image processing and Artia-Wrapper sub-tomogram averaging

The movie stacks were aligned to compensate for beam-induced movement using the MotionCor2 wrapper within RELION-5 (8), and the contrast transfer function subsequently estimated using CTFFIND4 v4.1.1426. The tilt series were then aligned using IMOD v4.11.24 patch tracking (9). The tomographic reconstructions were performed by various algorithms such as simultaneous algebraic reconstruction technique for high-contrast and weighted back-projection for sub-tomogram averaging (10).

Tomogram segmentation was performed using Dragonfly (v. 2024.1 for Windows), and the smoothness of the segmentations was improved using mean curvature motion (11) (https://github.com/FrangakisLab/mcm-cryoet). MATLAB scripts were used to measure the distance between two adjacent podocyte membranes and the periodicity of the densities forming the SD by cross-correlation (MATLAB 2023a, 2023 and MATLAB 2024a, 2024; The MathWorks, Natick, MA, USA, https://github.com/FrangakisLab/membraneDistanceMeasurement).

In the tomographic reconstructions, oblique planes were placed on the SD based on orientation of the glomerular basement membrane, and points on individual strands were selected on the plane using ArtiaX (12) implemented in UCSF ChimeraX (13). A total of 62 sub-tomograms (box size 128 pix, pixel size 5.36 Å) were extracted. All sub-tomograms showed a fishnet pattern but with a strong missing wedge, leading to a very strong signal anisotropy. Consequently, sub-tomogram averaging was used more to confirm the similarity of the data to the murine and nephrocyte models than for structural purposes.

#### Slit diaphragm modelling of the Nephrin-Neph1 heterodimers

The molecular model of Nephrin-Neph1 heterodimers was generated from predictions available from the AlphaFold Protein Structure Database (14, 15) (IDs: AF-Q9QZS7-F1, AF-Q80W68-F1) with the help of Coot, since no full-length structure is available to date. The unstructured regions at the N- and C-termini with low predicted confidence were removed (Nephrin: amino acids 1–29 and 1038–1241; Neph1: 1–19 and 490–757). The angle between Nephrin and Neph1 was estimated from the cryo-ET map at 90°. Nephrin and Neph1 were then assembled into a heterodimer based on the crystal structure of the SYG-2-SYG-1 heterodimer (ID: 4OFY) (5). The heterodimers were then placed according to the densities of the cryo-ET map. Prior information about their assembly was used from the murine SD, as the resolution from the human SD alone was insufficient for a reliable fit. The arrangement of the heterodimers was selected so that no unoccupied densities remained. The spacings of the heterodimers between each other as well as to the membrane were according to the measurement of the cryo-ET map.

#### Sex as a biological variable

This study, although involving a sample obtained from a male patient, did not consider sex as a biological variable.

#### Statistics

62 SD segments were combined to produce the final sub-tomogram average.

## Study approval

### Ethics approval and consent to participate

Tissue samples used in this study were provided by the University Hospital Frankfurt. Written informed consent was obtained from the patient, and the study was approved by the institutional Review Boards of the UCT and the Ethical Committee at the University Hospital Frankfurt (project-number: SUG-6-2018).

### Data availability

The cryo-ET data sets will be deposited to EMPIAR and to the Cryo-ET Data Portal.

## Supplementary figures

**Supplementary Figure S1:**
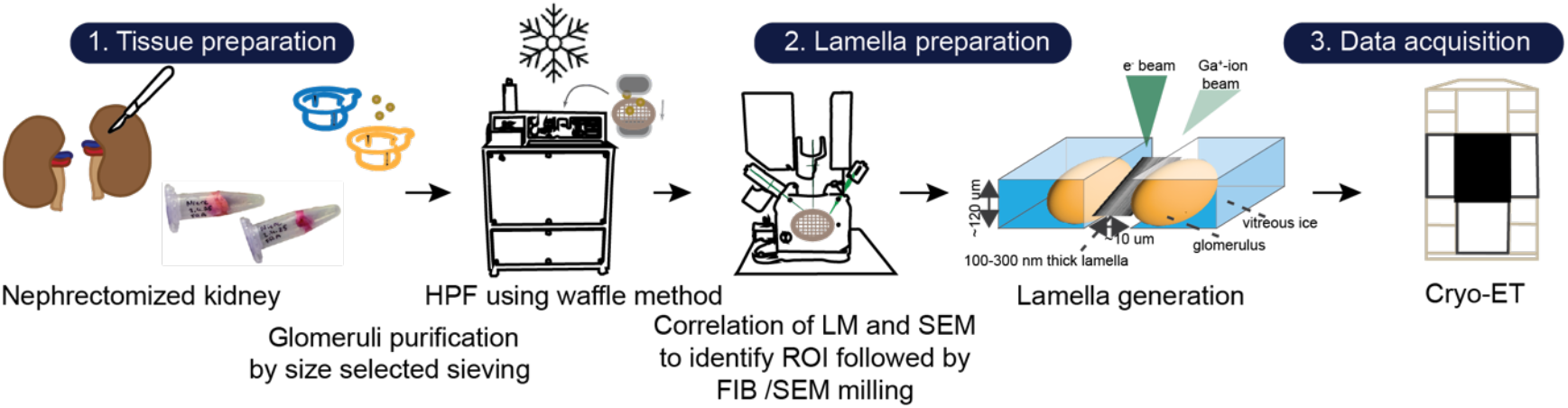
Glomeruli preparation workflow. Glomeruli were isolated from a nephrectomized human tissue by size-selected sieving and subjected to high-pressure freezing (HPF) to liquid nitrogen temperatures. A random area of high-pressure frozen sample was thinned to lamella of a thickness of ∼250 nm with a focused-ion beam (FIB) scanning electron microscope (SEM). Finally, the lamellae were imaged using cryo-electron tomography (cryo-ET).

**Supplementary Figure S2:**
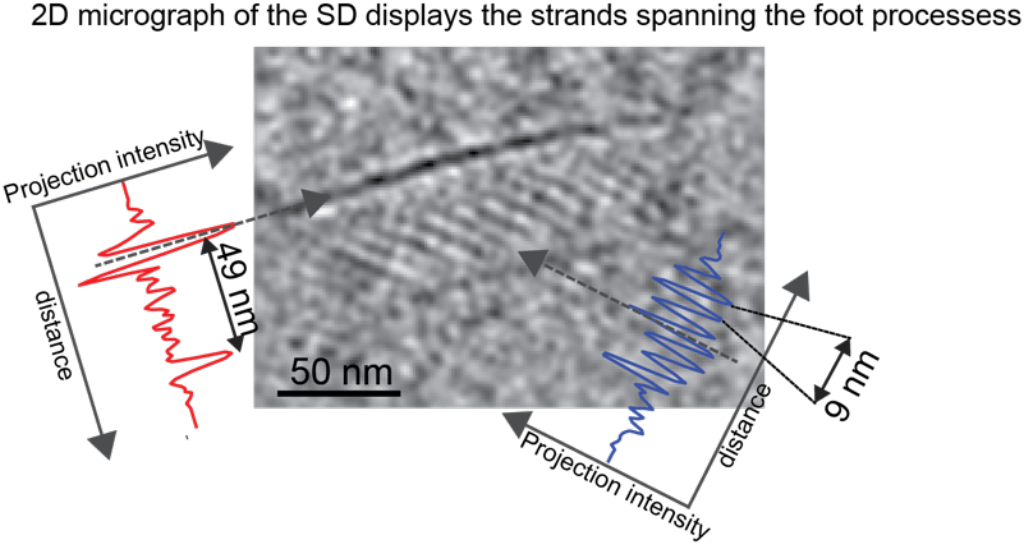
Example of a 2D electron micrograph with a particularly strong contrast. The strands spanning the two foot processes can be seen at a spacing of ∼9 nm. Because it is a 2D projection, only one layer of molecules is visible. The plots in the two directions display the spacing of the plasma membranes (in red) and the spacing of the strands (in blue).

**Supplementary Figure S3:**
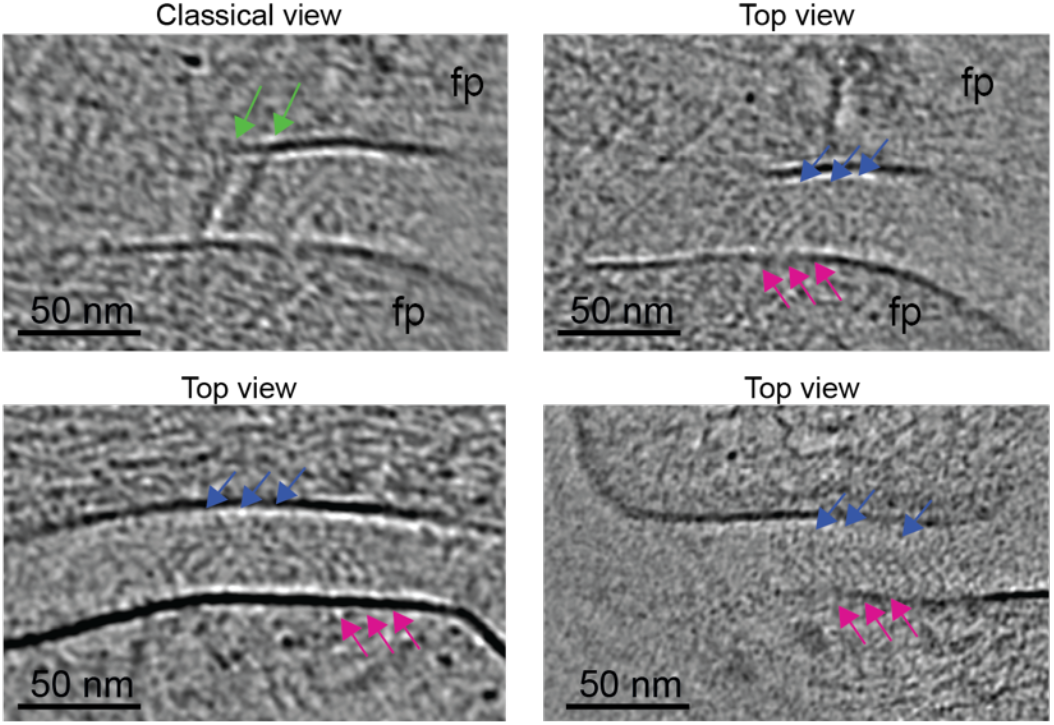
Four example of SDs seen in the cryo-electron tomograms. Computational sections through the SDs are shown (similar view to Figure 1c). The fishnet pattern with identical dimensions is visible in the noisy raw data. This signal was used for averaging and consequently improving the signal-to-noise ratio.

**Supplementary Figure S4:**
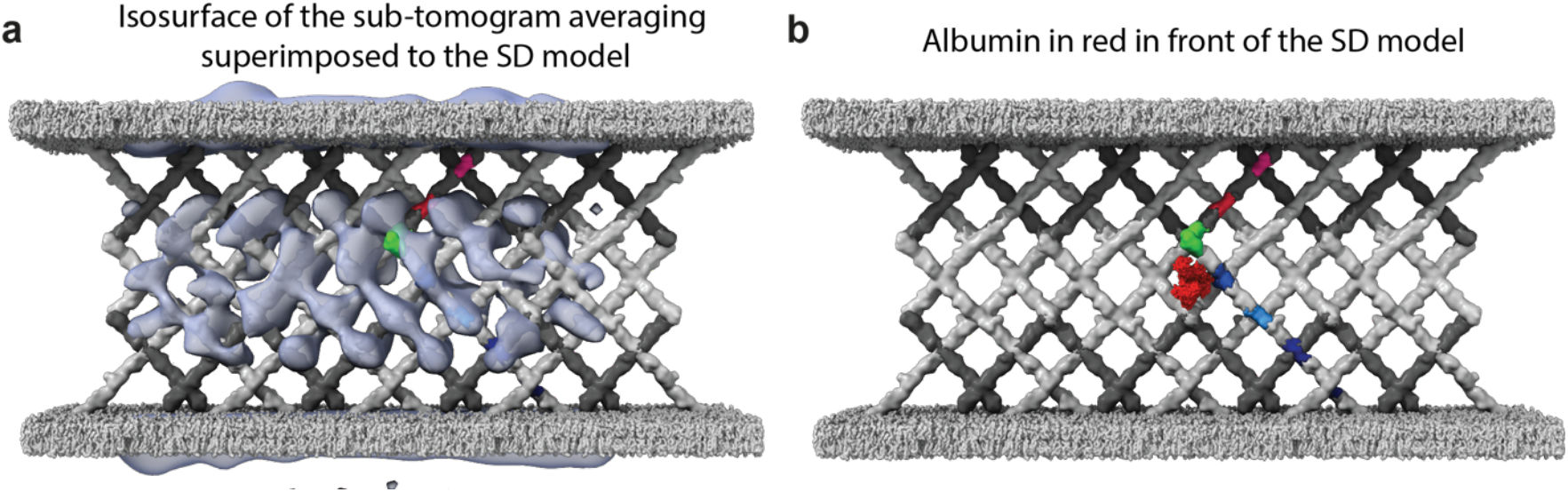
Models of the SD. (a) The SD model superimposed on the isosurface of the sub-tomogram average (in transparent blue). One heterodimer is shown in different colors highlighting the crossing points. (b) Superposition of albumin shown in red (PDB:1AO6) in front of the schematic of the SD, showing their relative dimensions. The size of the structural holes compared to albumin can be appreciated.

